# Hydrogen peroxide production by *Streptococcus pneumoniae* results in alpha-hemolysis by oxidation of oxy-hemoglobin to met-hemoglobin

**DOI:** 10.1101/2020.07.23.218966

**Authors:** Erin McDevitt, Faidad Khan, Anna Scasny, Zehava Eichembaun, Larry S. McDaniel, Jorge E. Vidal

## Abstract

*Streptococcus pneumoniae* (Spn) and other streptococci produce a greenish halo on blood agar plates referred to as α-hemolysis. This phenotype is utilized by clinical microbiology laboratories to report culture findings of α-hemolytic streptococci, including Spn, and other bacteria. The α-hemolysis halo on blood agar plates has been related to the hemolytic activity of pneumococcal pneumolysin (Ply), or to a lesser extent, to lysis of erythrocytes by Spn-produced hydrogen peroxide. We investigated the molecular basis of the α-hemolysis halo produced by Spn. Wild-type strains TIGR4, D39, R6, and EF3030, and isogenic derivative Δ*ply* mutants, produced a similar α-hemolytic halo on blood agar plates while cultures of hydrogen peroxide knockout Δ*spxB*/Δ*lctO* mutants lacked this characteristic halo. Spectroscopic studies demonstrated that culture supernatants of TIGR4 released hemoglobin-bound heme (heme-hemoglobin) from erythrocytes and oxidized oxy-hemoglobin to met-hemoglobin within 30 min of incubation. As expected, given Ply hemolytic activity, and that hydrogen peroxide contributes to the release of Ply, TIGR4 isogenic mutants Δ*ply* and Δ*spxB*/Δ*lctO* had a significantly decreased release of heme-hemoglobin from erythrocytes. However, TIGR4Δ*ply* that produces hydrogen peroxide oxidized oxy-hemoglobin to met-hemoglobin, whereas TIGR4Δ*spxB/*Δ*lctO* failed to produce oxidation of oxy-hemoglobin. We demonstrated that the so-called α-hemolysis halo is caused by the oxidation oxy-hemoglobin (Fe^+2^) to a non-oxygen binding met-hemoglobin (Fe^+3^) by Spn-produced hydrogen peroxide. Since Spn colonizes the human lung, oxidation of oxy-hemoglobin might have important implications for pathogenesis.

**Importance:** There is a misconception that α-hemolysis observed on blood agar plates cultures of *Streptococcus pneumoniae* (Spn), and other α-hemolytic streptococci is produced by a hemolysin, or alternatively, by lysis of erythrocytes caused by hydrogen peroxide. We noticed in the course of our investigations that wild-type Spn strains and hemolysin (e.g., pneumolysin) knockout mutants, produced the α-hemolytic halo on blood agar plates. In contrast, hydrogen peroxide defective mutants prepared in four different strains lacked the characteristic α-hemolysis halo. We also demonstrated that wild-type strains and pneumolysin mutants oxidized oxy-hemoglobin to met-hemoglobin. Hydrogen peroxide knockout mutants, however, failed to oxidize oxy-hemoglobin. Therefore, the greenish halo formed on cultures of Spn and other so-called α-hemolytic streptococci is caused by the oxidation of oxy-hemoglobin produced by hydrogen peroxide. Oxidation of oxy-hemoglobin to the non-binding oxygen form, met-hemoglobin, might occur in the lungs during pneumococcal pneumonia.

## Manuscript

Historically *S. pneumoniae* (Spn), and other streptococci of the viridians group, are classified as alpha (α)-hemolytic bacteria on the basis of a greenish halo that surrounds colonies when grown aerobically on blood agar plates (1, 2). This α-hemolytic activity has been related to the production of a hemolysin, which in the case of *S. pneumoniae* strains is referred as to pneumolysin, or Ply (3). Since anaerobic cultures of α-hemolytic streptococci lack this greenish discoloration, the phenotype has been also thought to be caused by lysis of erythrocytes by hydrogen peroxide produced as a byproduct of the metabolism of α-hemolytic streptococci (1). In this study we investigated whether Ply, or Spn-produced hydrogen peroxide, was responsible for α-hemolysis and identified the molecular basis of the phenotype.

The pneumolysin is encoded by *ply* (4), while hydrogen peroxide released in cultures of Spn strains and α-hemolytic streptococci is a byproduct of the metabolism of enzymes pyruvate oxidase (SpxB) and lactate dehydrogenase (LctO) (5). We utilized two different D39Δ*ply* mutants, TIGR4Δ*ply*, and *TIGR4*Δ*spxB*/Δ*lctO* from previous publications (6–8), and prepared Δ*spxB*/Δ*lctO* double mutants in three other backgrounds, vaccine serotype 19F strain EF3030 (9, 10), vaccine-escape serotype 2 strain D39, and R6 (11). We previously demonstrated by Western blot that the Ply knockout mutants do not produce Ply (6), and that *TIGR4*Δ*spxB*/Δ*lctO* does not produce detectable levels of hydrogen peroxide in supernatants from THY broth cultures incubated for 4 h (7).

Spn strains were inoculated on blood agar plates containing 5% sheep blood and plates were incubated at 37°C under aerobic conditions and a 5% CO_2_ atmosphere. Overnight cultures of TIGR4, D39, R6 (not shown), and EF3030 showed the classic α-hemolytic halo surrounding colonies (Fig. 1). The median diameter of α-hemolysis halos produced by D39 was 2.42 mm (Fig. 1, inset). Blood agar plates with cultures of isogenic TIGR4Δ*ply*, and D39Δ*ply*, mutant strains showed an undistinguishable α-hemolysis halo, with D39Δ*ply* producing α-hemolysis halos with a diameter of 2.26 mm (Fig. 1, inset). An additional TIGR4Δ*ply* mutant (AC4037) yielded a similar α-hemolysis halo [not shown, (12)]. In contrast, blood agar plates inoculated with TIGR4Δ*spxB*/Δ*lctO* and three additional double Δ*spxB*/Δ*lctO* mutants *R6*Δ*spxB*/Δ*lctO*, *D39*Δ*spxB*/Δ*lctO*, and *EF3030*Δ*spxB*/Δ*lctO*, completely lacked the α-hemolytic halo (Fig. 1). Blood agar plates of *TIGR4*Δ*spxB*/Δ*lctO*, *D39*Δ*spxB*/Δ*lctO* and *EF3030*Δ*spxB*/Δ*lctO* did not produce the α-hemolytic halo even after 72 h of incubation (not shown).

**Fig. 1.**
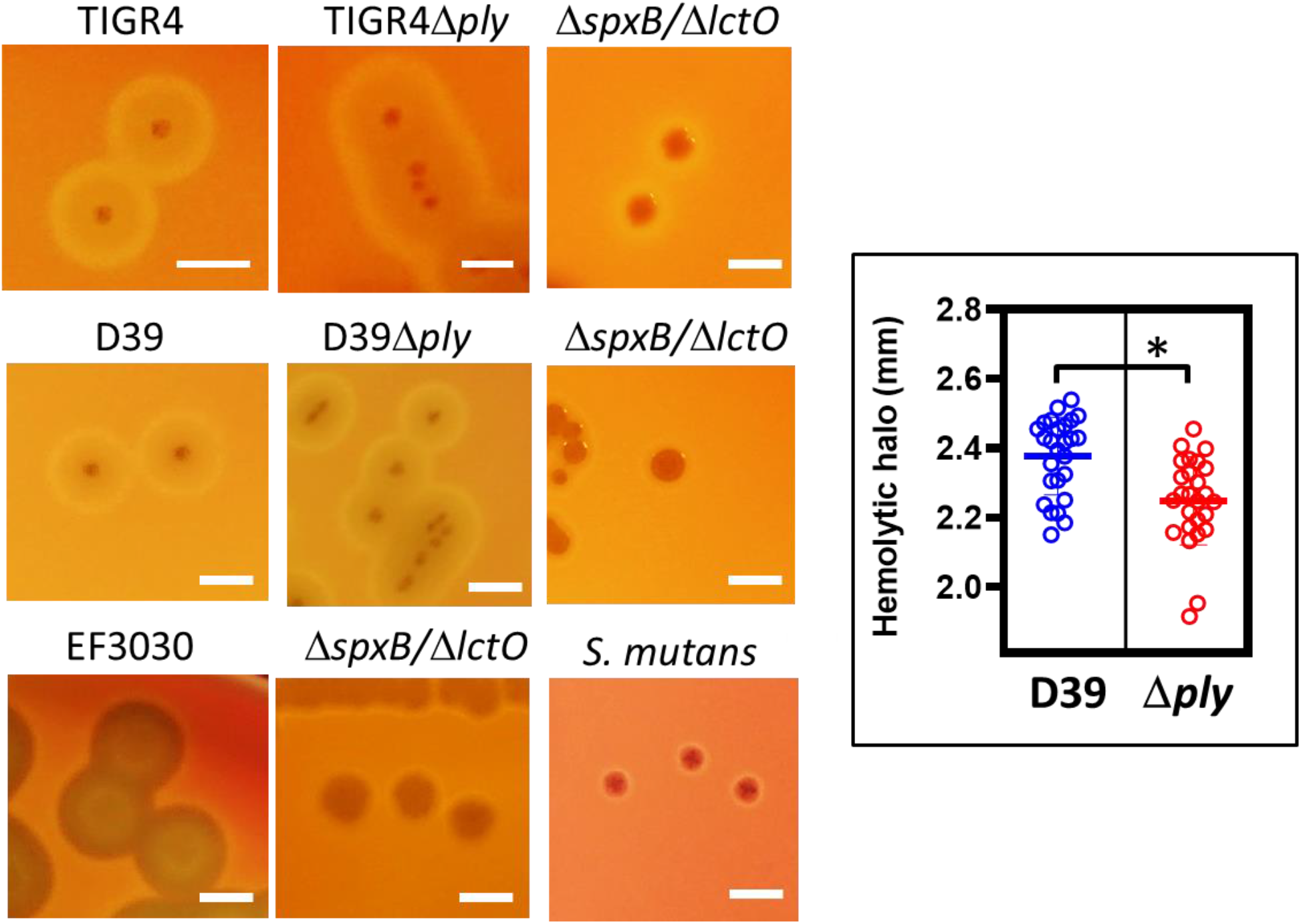
Hydrogen peroxide but not pneumolysin causes the α-hemolytic phenotype on blood agar plates. Spn wt strains TIGR4, D39, EF3030, or Δ*ply*, Δ*spxB*/Δ*lctO* mutant derivatives, or *S. mutans* strain ATCC 25175, was inoculated onto blood agar plates and incubated for 24 h at 37°C in a 5% CO_2_ atmosphere. Plates were photographed with a Canon Rebel EOS T5 camera system and digital pictures analyzed. Phenotypes were confirmed at least three times. Bars=2 mm. Inset: Hemolytic halo measured with ImageJ software on at least 25 colonies from images obtained from cultures on blood agar plates of D39 wt or *D39*Δ*ply*; unpaired Student *t* test was performed to assess significance; NS=no significant, *p*>0.05.

Other streptococci, including *S. mutans*, are not hemolytic when grown on blood agar plates (1, 2). Colonies of *S. mutans* strain ATCC 25175, on blood agar plates, resembled those of the *S. pneumoniae* isogenic hydrogen peroxide knockout mutants (Fig. 1). Similarly to *S. pneumoniae* and other streptococci, *S. mutans* encodes a putative α-hemolysin (13). *S. mutans* lacks production of detectable hydrogen peroxide in the supernatant when grown in aerobic cultures, and it is highly susceptible to hydrogen peroxide produced by α-hemolytic oral streptococci (14, 15). Together, this evidence indicates that hydrogen peroxide but not the hemolysin Ply caused the α-hemolytic phenotype observed in aerobic cultures of *S. pneumoniae* strains.

Erythrocytes carry hemoglobin that reversibly binds oxygen through a penta-coordinate heme molecule containing ferrous iron (Fe^+2^), known as oxy-hemoglobin (16). When hemoglobin is released from erythrocytes, heme-hemoglobin can be observed by optical spectroscopy at ~415 nm (16–18). This region is known as Soret region peak and represent heme-hemoglobin, while oxy-hemoglobin is characterized by two absorption peaks of ~540 and ~570 nm (17, 18). Oxy-hemoglobin (Fe^+2^) is autoxidized to met-hemoglobin (Fe^+3^), or oxidized by radicals such as hydrogen peroxide (17, 18), inducing spectral changes, i.e., flattening the oxy-hemoglobin absorbance peaks. Spn produces and releases abundant hydrogen peroxide into the culture supernatant that intoxicate human cells (19), or that rapidly kills *Staphylococcus aureus* strains and other bacterial species (7, 20). Hydrogen peroxide is a byproduct of the metabolism of two different enzymes, pyruvate oxidase (SpxB) and lactate dehydrogenase (LctO) (5).

To further confirm that the α-hemolytic phenotype of Spn was due to hydrogen peroxide, we utilized a modified hemoglobin release assay that, when coupled with optical spectroscopy, allowed us to quantify the release of heme-hemoglobin and to observe the oxidation of oxy-hemoglobin to met-hemoglobin. As a control of heme-hemoglobin release and the presence of oxy-hemoglobin, we obtained the UV-visible absorption spectra of a 3% suspension of sheep erythrocytes that had been lysed with an equal volume of water, or lysed with 0.1% final concentration of saponin (Fig. 2A and not shown). After centrifugation of the lysed erythrocytes suspension at 300 x *g* for 5 min no red blood cells were visible in the bottom, and therefore this was considered as the maximum heme-hemoglobin released. As expected three characteristic peaks were observed. The Soret peak, which wavelength of maximum absorption was 415 nm, and its absorbance was set as to 100% hemoglobin release (Fig. 2A), and two oxy-hemoglobin peaks at 540 and 570 nm (Fig. 2A). Similar peaks were observed when hemoglobin was released from sheep, or horse, erythrocytes with saponin (not shown). To investigate release of heme-hemoglobin, Spn strains were inoculated in THY broth (pH=7) and incubated at 37°C in a 5% CO_2_ atmosphere for 1, 2, 3, or 4 h. Bacteria-free supernatants were harvested, and then incubated with an equal volume of a 3% suspension of sheep erythrocyte at 37°C in a 5% CO_2_ atmosphere for 30 min, after which the treated erythrocytes suspensions were centrifuged at 300 x *g* for 5 min to collect supernatants.

**Fig. 2.**
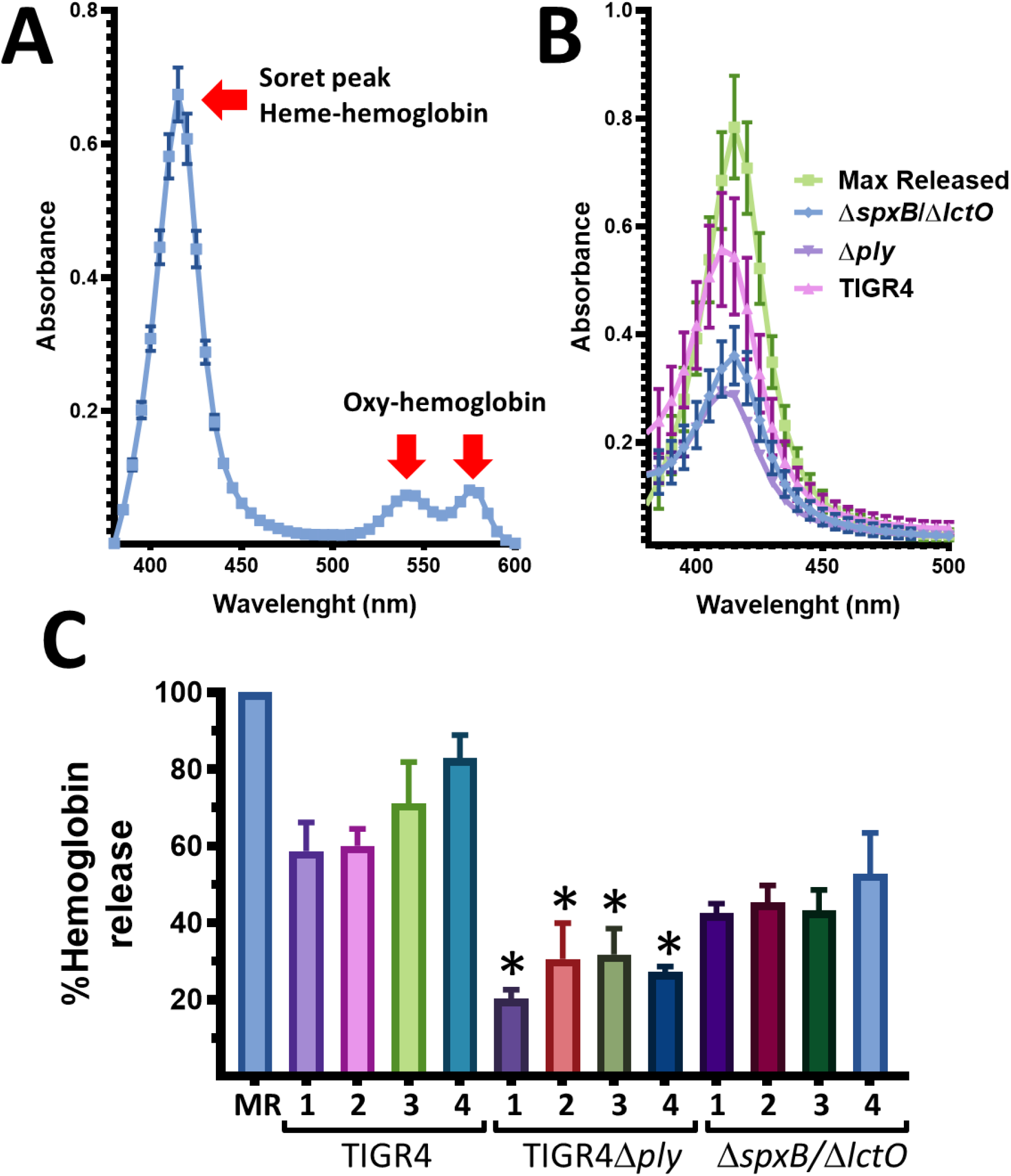
Heme-hemoglobin release by Spn strains. (A) Suspension (3%) of sheep erythrocytes was lysed, centrifuged at 300 x *g* for 5 min and, after incubation for 30 min at 37°C in a 5% CO_2_ atmosphere, the spectra was obtained using an spectrophotometer Omega BMG LabTech (ThermoFisher). (B and C) TIGR4 wt, TIGR4Δ*ply*, or TIGR4Δ*spxB/*Δ*lctO*, was inoculated in THY broth (pH=7.0) and incubated during 4 h at 37°C in a 5% CO_2_ atmosphere. Bacteria-free supernatants were harvested by centrifugation at 13,000 x *g* for 5 min and an equal volume was incubated with a 3% suspension of sheep erythrocytes for 30 min at 37°C. After pelleting down the erythrocytes as above, the hemoglobin-containing supernatant was collected and the UV-visible absorption spectrum was obtained. Panel (B) shows the 4 h time point. In panel (C), maximum heme-hemoglobin release was set to 100% and the % release by culture supernatants was calculated. Error bars represent the standard errors of the means calculated using data from at least three independent experiments.*p<0.05 comparing Soret absorbance at 415 generated by wt and isogenic mutant, at the same time point.

A time course study demonstrated that TIGR4 released ~60% of heme-hemoglobin as soon as 1 h post-inoculation (Fig. 2C) and produced, after 4 h of incubation, a Soret peak with a comparable absorbance to that of the heme-hemoglobin released in the control (Fig. 2B) that represented ~85% of heme-hemoglobin released compared to the maximum heme-hemoglobin released control (Fig. 2C). As expected given that hydrogen peroxide contributes to release of Ply into the supernatant (21), hemoglobin released by *TIGR4*Δ*spxB*/Δ*lctO* after 4 h of incubation was ~50% of that released by the wt strain (Fig. 2B and 2C). In contrast to the TIGR4 strain and *TIGR4*Δ*spxB*/Δ*lctO*, the isogenic Ply knockout mutant TIGR4Δ*ply* induced the release of <30% heme-hemoglobin compared to the control, indicating that either another hemolysin, or hydrogen peroxide produced by this Ply mutant bear some hemolytic activity against sheep erythrocytes. These results support the hypothesis that the α-hemolytic phenotype observed on blood agar plates is not due to Ply-associated hemolytic activity but to hydrogen peroxide produced by the pneumococcus.

The presence of oxy-hemoglobin peaks, which were clearly observed in the control preparation (Figs. 2A and 3A), were completely flattened when culture supernatants of TIGR4 strain obtained after 3, or 4 h of incubation, and were incubated with the suspension of erythrocytes for an additional 30 min period. This change in the absorption pattern of oxy-hemoglobin was compatible with the oxidation of oxy-hemoglobin to met-hemoglobin, a non-oxygen binding form of hemoglobin (22). Because culture supernatants from TIGR4Δ*ply*, or TIGR4Δ*spxB/*Δ*lctO*, did not release heme-hemoglobin at the same level to that of the TIGR4, we could not evaluate the oxidation of oxy-hemoglobin in these isogenic mutant strains using the modified hemoglobin release assay.

**Fig. 3.**
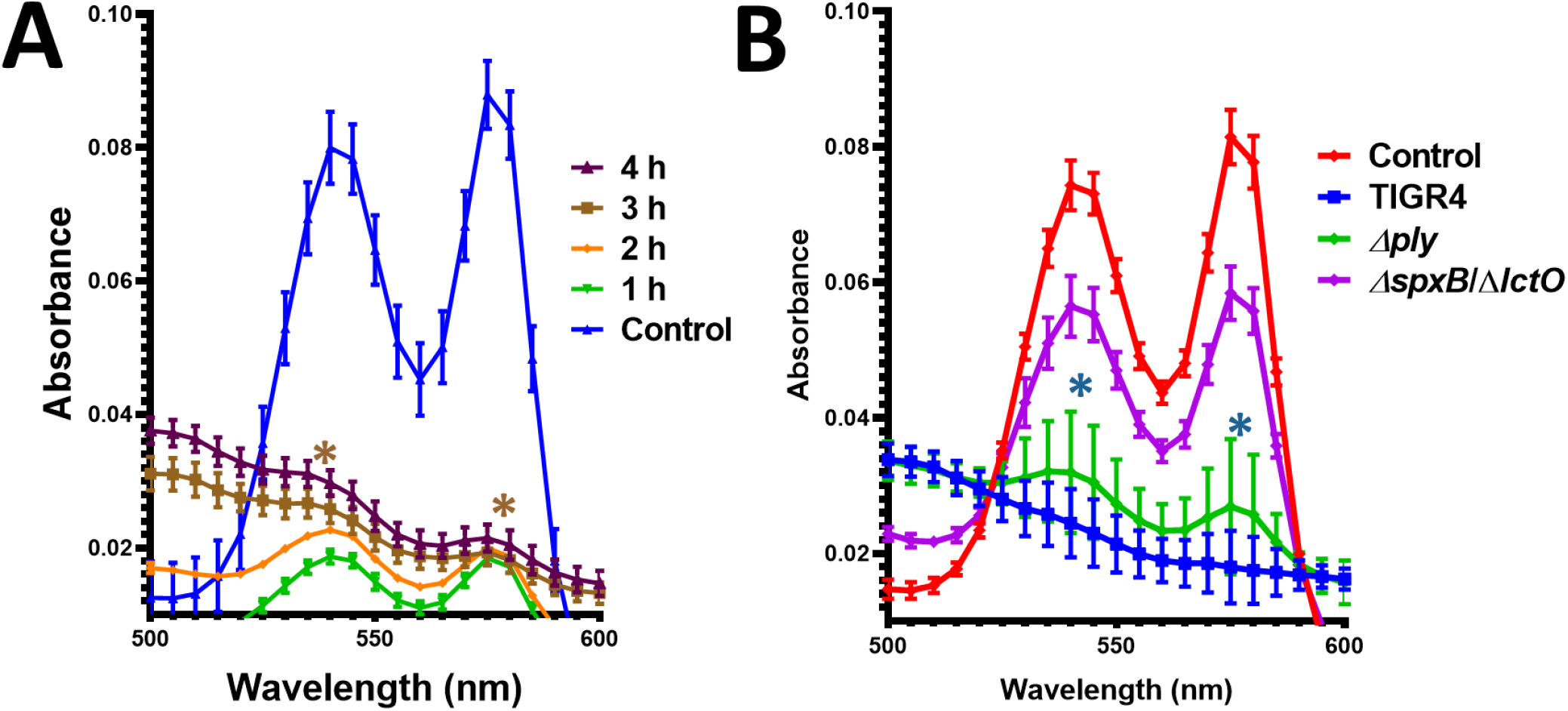
Oxy-hemoglobin is oxidized to met-hemoglobin by Spn-produced hydrogen peroxide. (A) Suspension (3%) of sheep erythrocytes was mixed with an equal volume of cell-free culture supernatants of strain TIGR4 wt that had been grown as in Fig. 2B for the indicated time. The mixture was incubated for 30 min at 37°C in a 5% CO_2_ atmosphere. As a control, erythrocytes were lysed and incubated under the same conditions. The absorbance spectra was then obtained using an spectrophotometer Omega BMG LabTech (ThermoFisher). (B) TIGR4 wt, TIGR4Δ*ply*, or TIGR4Δ*spxB/*Δ*lctO*, was inoculated in THY broth (pH=7.0) and incubated during 4 h at 37°C in a 5% CO_2_ atmosphere. Bacteria-free supernatants were harvested by centrifugation at 13,000 x *g* for 5 min and an equal volume was incubated for 30 min at 37°C with hemoglobin-containing erythrocytes lysates. Hemoglobin-containing supernatants were collected and the UV-visible absorption spectrum was obtained. Error bars represent the standard errors of the means calculated using data from at least three independent experiments.*p<0.05 comparing the oxy-hemoglobin absorbance peaks (540 nm and 570 nm) generated by the untreated hemoglobin-containing lysate control against that generated by incubation with supernatants from TIGR4 wt or TIGR4Δ*ply* strain.

To further confirm whether oxy-hemoglobin is oxidized to met-hemoglobin by hydrogen peroxide produced in culture supernatants of TIGR4Δ*ply* but not in supernatants of hydrogen peroxide knockout mutant TIGR4Δ*spxB/*Δ*lctO*, we incubated preparations of oxy-hemoglobin that had been previously released from erythrocytes, as mentioned earlier, with culture supernatants of TIGR4 or isogenic mutants. We reason that if hydrogen peroxide present in culture supernatants was responsible for the oxidation of oxy-hemoglobin, then having oxy-hemoglobin already as a substrate would allow us to observe such a reaction. As expected, supernatants from 4 h cultures of the wt strain that were incubated for 30 min with the oxy-hemoglobin preparation, converted oxy-hemoglobin to met-hemoglobin (Fig. 3B). Supernatants from the isogenic TIGR4Δ*ply* significantly oxidized oxy-hemoglobin to met-hemoglobin indicating that oxidation occurred due to the hydrogen peroxide activity retained by the *ply* knockout mutant. Confirming this hypothesis, oxy-hemoglobin was observed almost intact after 30 min incubation with supernatants of the isogenic *TIGR4*Δ*spxB*/Δ*lctO*, indicating that met-hemoglobin was produced by hydrogen peroxide secreted into the culture supernatant. A similar oxidation of hemoglobin to met-hemoglobin was observed using horse erythrocytes, and releasing hemoglobin from erythrocytes using water or saponin (not shown).

In conclusion, we demonstrated in this study that the so-called α-hemolysis phenoype observed on blood agar plates, when incubated under aerobic conditions, is an oxidative reaction cause by Spn-produced hydrogen peroxide that converts oxy-hemoglobin to met-hemoglobin.

## Acknowledgements

This study was supported in part by a grant from the National Institutes of Health (NIH; 1R21AI144571-01 to JEV) and generous start-up funds from the University of Mississippi Medical Center (UMMC). The content is solely the responsibility of the authors and does not necessarily represent the official view of the NIH or UMMC. We thanks Dr. Andrew Camilli from Tufts University and Dr. James C. Paton from University of Adelaide for the kind gift of strain AC4037 and D39Δ*ply*, respectively. We would also like to express our gratitude to Dr. Bernard Beall and Dr. Lesley McGee from the Centers for Disease Control and Prevention for providing *S. mutans*. Authors thanks Professor David Stephens and Emilio Rodriguez from Emory University School of Medicine, for earlier discussions of preliminary findings, and for his assistance on some procedures, respectively.

